# FOP plays a negative role in ciliogenesis and promotes cell cycle re-entry by facilitating primary cilia disassembly

**DOI:** 10.1101/627349

**Authors:** Huadong Jiang, Man-Hei Cheung, Aftab Amin, Chun Liang

**Affiliations:** Division of Life Science, Center for Cancer Research and State Key Lab for Molecular Neuroscience, Hong Kong University of Science and Technology, Hong Kong, China; Guangzhou HKUST Fok Ying Tung Research Institute, Guangzhou, China; Intelgen Limited, Hong Kong-Guangzhou-Foshan, China

**Keywords:** FOP, primary cilia, cilia assembly and disassembly, cell cycle exit and re-entry, actin dynamics

## Abstract

Primary cilia are microtubule-based, antenna-like organelles, which are formed in G_0_ phase and resorbed as cells re-enter the cell cycle. It has been reported that the length of primary cilia can influence the timing of cell cycle progression. However, the molecular links between ciliogenesis and cell cycle progression are not clear. FOP (Fibroblast growth factor receptor 1 Oncogene Partner, also known as FGFR1OP) has been implicated in ciliogenesis. Here, we show that the expression of FOP during cell cycle exit and re-entry is negatively correlated with ciliogenesis. Knockdown of FOP promotes cilia elongation and suppresses timely cilia disassembly. In contrast, ectopic expression of FOP inhibits cilia growth. Moreover, pharmacological inhibition of actin polymerization with Cytochalasin D abrogates FOP-induced cilia disassembly, suggesting that FOP facilitates cilia disassembly by promoting actin cytoskeleton formation. Lastly, knockdown of FOP delays cell cycle re-entry of quiescent cells following serum re-stimulation, and this can be reversed by silencing IFT20 (intraflagellar transport 20), an intraflagellar transport protein essential for ciliogenesis. Collectively, these results suggest that FOP plays a negative role in ciliogenesis and can promote cell cycle re-entry by facilitating cilia disassembly.

## Introduction

Primary cilia are microtubule-based organelles that protrude from the cell apical surface to sense environmental cues that regulate cell growth, development and homeostasis (Ishikawa and Marshall, 2011; Malicki and Johnson, 2017; Nigg and Raff, 2009; Satir and Christensen, 2007; Satir et al., 2010). As cell signaling centers, primary cilia coordinate with many cell signaling pathways, including those mediated by Hedgehog, Wnt and RTKs (Receptors of Tyrosine Kinases) (Bangs and Anderson, 2017; Christensen et al., 2017; Liu et al., 2018; Malicki and Johnson, 2017; Wallingford and Mitchell, 2011). Defects in cilia assembly and/or functions can lead to a wide range of human disorders, termed ciliopathies, characterized by mental retardation, polycystic kidney, retinal defects, obesity, diabetes and other development abnormalities (Hildebrandt et al., 2011; Reiter and Leroux, 2017).

The assembly and disassembly of primary cilia are tightly controlled in the cell cycle. Primary cilia are formed in quiescent (G_0_ phase) cells and are resorbed as cells re-enter the cell cycle (Sánchez and Dynlacht, 2016). Primary cilia emanate from the mother centrioles, and primary cilia and centrosomes share the same centrioles. When a quiescent cell enters the proliferative cycle, the centrioles need be released from primary cilia in order to act as spindle poles in mitosis. It is therefore assumed that primary cilia function as a structural checkpoint for cell cycle re-entry (Basten and Giles, 2013; Izawa et al., 2015; Ke and Yang, 2014). This hypothesis has been verified by several studies. For example, both Nde1 (Nuclear distribution gene homologue 1) and Tctex-1 (dynein light chain tctex-type 1, also known as DYNLT1) promote primary cilia disassembly and accelerate cell cycle re-entry in response to growth-factor stimulation, and they do not significantly influence cilia assembly (Kim et al., 2011; Li et al., 2011). Moreover, knockdown of NEK24 or KIF24 induces primary cilia formation and inhibits cell proliferation in cycling RPE1 cells and breast cancer cells (Kim et al., 2015). In contrast, depletion of proteins required for ciliogenesis, such as IFT88 (intraflagellar transport 88, also known as Polaris and Tg737) and KIF3A (kinesin family member 3A), facilitates cell proliferation (Deng et al., 2018).

The dynamics of primary cilia assembly and disassembly is modulated by numerous regulators (Sánchez and Dynlacht, 2016). Recent studies have shown that the actin cytoskeleton is involved in this dynamic process. Pharmacological inhibition of actin polymerization with Cytochalasin D (CytoD) promotes primary cilia formation and elongation and inhibits cilia disassembly during cell cycle re-entry (Kim et al., 2011; Li et al., 2011; Sharma et al., 2011), suggesting that the actin cytoskeleton negatively regulates ciliogenesis. Moreover, actin related proteins such as cortactin, components of the ARP2/3 complex and Tctex-1 required for actin nucleation, branching, polymerization and stress fiber formation, also inhibit ciliogenesis and promote primary cilia disassembly (Bershteyn et al., 2010; Cao et al., 2012; Drummond et al., 2018; Kim et al., 2010; Ran et al., 2015; Saito et al., 2017).

While primary cilia assembly has been extensively studied, much less is known about the molecular mechanism underlying primary cilia disassembly (resorption). In addition to the proteins mentioned above, several other signaling axes, such as Aurora A-HDAC6 (Histone Deacetylase 6), responsible for cilia disassembly have been identified (Sánchez and Dynlacht, 2016). Aurora A can be activated by HEF1 (Human enhancer of filamentation 1, also known as NEDD9), Pitchfork (Pifo), trichoplein and calcium/calmodulin, and the activated Aurora A in turn phosphorylates HDAC6 and stimulates its tubulin deacetylation activity, resulting in the destabilization of the ciliary axoneme and thus cilia resorption (Inoko et al., 2012; Kinzel et al., 2010; Plotnikova et al., 2012; Pugacheva et al., 2007).

FOP (Fibroblast growth factor receptor Oncogene Partner, also known as FGFR1OP) is a centrosomal and centriolar satellite protein (Lee and Stearns, 2013; Popovici et al., 1999; Yan et al., 2006). It associates with CEP350 and is required for microtubule anchoring (Yan et al., 2006). Knockout of FOP in chicken DT40 lymphocytes causes G_1_ arrest and eventual apoptosis (Acquaviva et al., 2009). Although several studies have suggested that FOP is involved on ciliogenesis, the role of FOP on ciliogenesis remains controversial. Some studies reported that knockout of FOP inhibited primary cilia formation (Kanie et al., 2017; Lee and Stearns, 2013; Mojarad et al., 2017); however, moderate knockdown of FOP had no effect on ciliogenesis (Graser et al., 2007; Lee and Stearns, 2013). Moreover, whether FOP regulates cilia length is unknown. In this study, in contrast to previous reports, we show that FOP plays a negative role in ciliogenesis. Knockdown of FOP increases cilia length, while ectopic over-expression of FOP suppresses cilia growth. Moreover, FOP induces actin polymerization-dependent cilia disassembly. We also demonstrate that knockdown of FOP delays cell cycle re-entry of quiescent cells following serum re-stimulation. Disruption of ciliogenesis by IFT20 depletion abolishes the delay in cell cycle re-entry caused by FOP knockdown. Together, these data suggest a negative effect of FOP on ciliogenesis and reveal a novel role of FOP in cilia disassembly by promoting actin cytoskeleton formation and thereby linking the dynamics of ciliogenesis to cell cycle progression.

## Materials and Methods

### Cell Line and Culture

hTERT-RPE1 cells were obtained from the American Type Culture Collection (ATCC) and cultured in DMEM/F12 medium (Life Technologies) supplemented with 10% fetal bovine serum (FBS) and 0.01 mg/mL hygromycin B in a humidified incubator at 37 °C with 5% CO_2_.

### Plasmid Construction, Stable and Transient Transfection

GFP-tagged human FOP expression plasmid for mammalian cells was constructed by PCR and standard cloning techniques. Briefly, the human FOP (NM_007045.3) open reading frame (ORF) was amplified from human cDNA (reverse transcription products from total RNA isolated from HEK293T cells) using the following primers: FOP/Forward: 5’-CGGAATTCCGAGCAAGATGGCGGCGAC-3’, FOP/Reverse: 5’-GGGGTACCCCTGCAACATCTTCCAGATAATC-3’.

The PCR product was purified, cut with EcoR I and Xho I and then inserted into pEGFP-N1 (Clontech). The construct was verified by DNA sequencing.

Cell transfection with plasmid DNA was performed using Lipofectamine 2000 (Invitrogen) according to the manufacturer’s instructions. RPE1 cells were plated 12-24 hrs before transfection. The cell confluency was about 80% at the time of transfection. For the generation of FOP-GFP stable cell lines, 48 hrs post transfection, RPE1 cells were selected with 2 mg/mL G418 for approximately 2 weeks. The expression level of FOP-GFP in individual clones were determined by immunoblotting.

Cell transfection with siRNAs was performed using Lipofectamine RNAiMAX (Invitrogen) according to the manufacturer’s instructions. Cells were seeded 12 hrs before transfection. The cell confluency was approximately 30% at the time of transfection. The final siRNAs concentrations were 40 nM for FOP and 80 nM for IFT20. For double transfection, RPE1 cells were first transfected with either the negative control siRNA or IFT20 siRNA. Cells were then split about 24 hrs after the first transfection. After another 24 hrs of culturing, cells were transfected with the negative control siRNA or FOP siRNA for 48 hrs. Pooled FOP siRNA (sense strands 5’-CCCAUUCCUAAGCCAGAGAAA-3’, 5’-CGAGAGAAUUUAGCCCGAGAU-3’, 5’-GGAUCACUUGGAUUAGGAA-3’, and 5’-GCCCGAGAUUUAGGUAUAA-3’), IFT20 siRNA (5’-GCUCGGAACUUGCUCAAAU-3’), or negative control siRNA (sense strand 5’-UUCUCCGAACGUGUCACGU-3’). All siRNAs were purchased from Genepharma (Shanghai, China).

### RNA Isolation, Reverse Transcription and Real-Time PCR

Total RNA was isolated from cultured cells using TRIzol Reagent (Invitrogen), according to the manufacturer’s instructions. Reverse transcription was performed using QuantiTect Reverse Transcription Kit (QIAGEN). Real-time PCR was performed using FastStart Universal SYBR Green Master (Rox) (Roche) and LightCycler384 (Roche). The following PCR primers were used: IFT20/F: 5’-AGCAGACCATAGAGCTGAAGG-3’, IFT20/R: 5’-AGCACCGATGGCCTGTAGT-3’, β-actin/F: 5’-TCCTTCCTGGGCATGGAGTCCT-3’, β-actin/R: 5’-TGCCAGGGCAGTGATCTCCT-3’.

### Immunoblotting

Cells were harvested, washed with PBS, and lysed in RIPA buffer (150 mM NaCl,50 mM Tris-HCl, 0.1% SDS,1% NP-40 and 1% Triton X-100) supplemented with 1 mM PMSF and protease inhibitor cocktail (Roche) at 4°C for 20 min. The lysates were then centrifuged for 15 min at 12,000 rpm at 4°C. The supernatants were collected, and an equal volume of 2XLaemmli’s buffer was added. The sample was denatured for 5 min at 95 °C. Proteins were resolved by 12.5% SDS-PAGE gel and then transferred to nitrocellulose membranes (Pall Corporation). Membranes were blocked with 5% non-fat milk in TBST (0.1% Tween 20) for 1 hr before incubation with primary and secondary antibodies. Signals were detected using SuperSignal West Pico Chemiluminescent Substrate (Thermo Scientific) according to the manufacturer’s instructions. The following antibodies were used: rabbit anti-FOP (Abcam, ab156013, 1:2000), rabbit anti-Aurora A (Cell signaling Technology, 14475, 1:2000), rabbit anti-p-Aurora A (Thr288) (Cell signaling Technology, 3079, 1:500), rabbit anti-cyclin A2 (Abcam, ab18159, 1:10000), rabbit anti-p-cdc2 (Tyr15) (Cell Signaling Technology, 9111, 1:2000), rabbit anti-p-Rb (Ser807/811) (Cell Signaling Technology, 8516, 1:2000), and mouse anti-β-actin (Sigma, A5441, 1: 5000).

### Immunofluorescence Staining

Cells were fixed with ice-cold methanol for 5 min at −20°C or 4% PFA for 15 min at room temperature, permeabilized with 0.5% Triton X-100 for 5 min and blocked in 5% BSA for 1 hr at room temperature. The following primary antibodies were used: rabbit anti-γ-tubulin (Sigma, T5192, 1:1000), mouse anti-γ-tubulin (Sigma, T5326, 1:1000), mouse anti-acetylated tubulin (Sigma, T6793, 1: 1000), rabbit anti-Arl13b (Proteintech, 17711-1-AP, 1:1000), and rabbit anti-OFD1 (Sigma, HPA031103, 1:200). Secondary antibodies used were goat anti-mouse Alexa 488 (Thermo Fisher Scientific, 1:500) and goat anti-rabbit Alexa 594 (Thermo Fisher Scientific, 1:500). F-actin was stained with Alexa Fluor 594 phalloidin (Thermo Fisher Scientific), according to the manufacturer’s instructions. The nuclei were stained with DAPI (Thermo Fisher Scientific). Mounted slides were observed and photographed using Elipse Ti-E microscopy using a 60× oil objective. Images were acquired with NIS-elements basis research (Nikon) and processed with Image J (NIH).

### Flow Cytometry (FACS Analysis)

Cells were detached with trypsin, centrifuged at 3000 rpm for 10 min, washed twice with PBS and fixed overnight in 70% ethanol at −20°C. The cells were then resuspended and stained with PI staining buffer (5 μg/mL RNase A, 0.1% (v/v) Triton X-100, 10 mM EDTA (pH8.0) and 50 μg/mL propidium iodide [Sigma]) for 1 hr at room temperature. For each sample, at least 10,000 cells were analyzed by FACSort machine (Becton Dickinson). Data were analyzed using Flowing Software 2.

### Cilia Assembly and Disassembly Assays

To induce primary cilia assembly, cells (untreated or 24 hrs post transfection) were starved in serum-free medium for 48 hrs. In some experiments, cells were treated with DMSO or 1 μM Cytochalasin D (CytoD, Sigma) at the final 2 hrs of the serum-starvation. To analyze cilia disassembly, cells were plated at less than 30% confluency before transfection. Cells were serum starved for 48 hrs immediately after transfection, and then 10% serum was added back to the medium to stimulate cilia resorption and cell cycle re-entry. Cells were fixed at different time points and immunostained with cilia makers. The length of the cilia was measured using Image J software (NIH).

### EdU Incorporation Assay

EdU (10 μM) was added to the growth medium 2 hrs prior to fixation. The cells were subsequently stained according to the manufacturer’s instructions (RiboBio). Mounted slides were observed and photographed using Elipse Ti-E microscopy using a 20X objective. Images were acquired with NIS-elements basis research (Nikon) and processed with Image J (NIH).

### Statistical Analysis

Statistical analysis was performed with Prism version 5 (GraphPad). Statistical significance of the difference between two groups was determined as indicated. Statistically significant differences were defined as *p<0.05, **p<0.01, and ***p<0.001.

## Results

### The FOP Expression Level Is Negatively Correlated with Ciliogenesis

Although several studies have suggested that FOP may be involved in ciliogenesis (Cabaud et al., 2018; Kanie et al., 2017; Mojarad et al., 2017), the regulation of FOP expression during cilia assembly and disassembly remains unclear. We first determined the correlation between the expression of FOP and ciliogenesis during cell cycle exit and re-entry, human telomerase-immortalized retinal pigmented epithelial (hTERT-RPE1, hereafter RPE1) cells were serum starved to induce primary cilia formation and cell cycle exit, and re-stimulated with serum to disassemble cilia and cell cycle re-entry. The protein expression levels of FOP were examined by immunoblotting, and the percentage of ciliated RPE1 cells were examined by immunostaining at different time points during serum starvation and re-stimulation. We found that the FOP expression level decreased and the percentage of ciliated cells increased upon serum starvation, while the FOP expression level increased and the percentage of ciliated cells decreased after serum re-stimulation (Figure 1A and 1B). The results indicate a negative correlation between FOP protein expression and the percentage of ciliated cells, suggesting a possible negative role of FOP in ciliogenesis.

**Figure 1.**
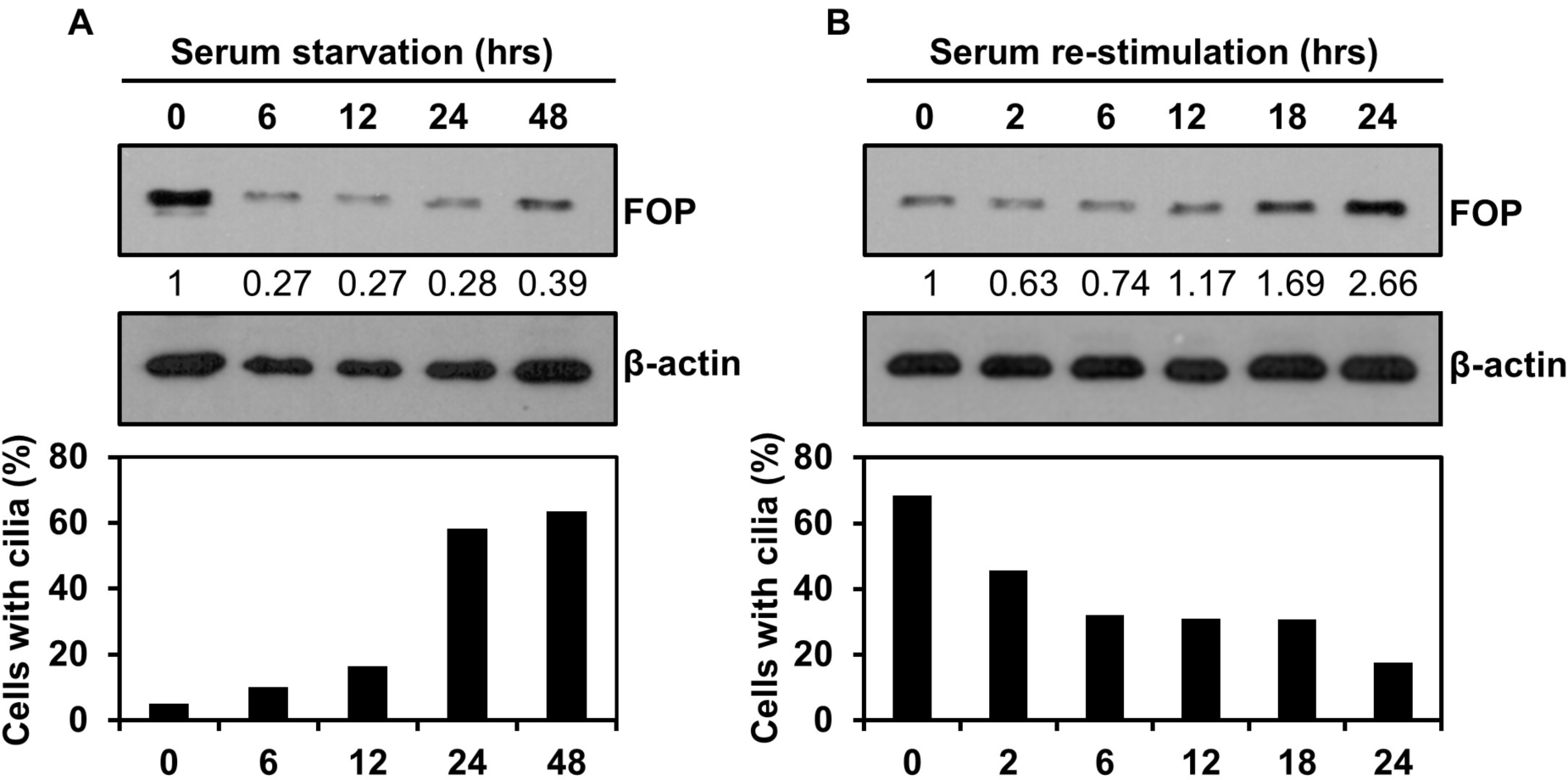
The FOP expression level negatively correlates with ciliogenesis. (**A**) The protein level of FOP decreases during cilia assembly. RPE1 cells were serum-starved for 0 hr, 6 hrs, 12 hrs, 24 hrs, and 48 hrs to induce primary cilia formation. The expression of FOP during cilia assembly were determined by immunoblotting (top) and the percentage of ciliated cells were determined by acetylated tubulin and *γ*-tubulin immunostaining (bottom; only the quantification is shown). (**B**) The protein level of FOP increases during cilia disassembly. After 48 hrs of serum starvation, RPE1 cells were re-stimulated with 10% serum for 0 hr, 2 hrs, 6 hrs, 12 hrs, 18 hrs and 24 hrs. The expression of FOP during cilia disassembly were determined by immunoblotting (top) and the percentage of ciliated cells were determined by acetylated tubulin and *γ*-tubulin immunostaining (bottom; only the quantification is shown). At least 200 cells were quantified for each time point.

### FOP Plays a Negative Role in Ciliogenesis

To confirm the negative role of FOP in ciliogenesis, we silenced FOP in RPE1 cells using siRNA (Figure 2A) and examined the effects on ciliogenesis by Arl13b (ADP-ribosylation like protein) immunostaining (Figure 2B-D). Surprisingly, the percentage of ciliated cells in the negative control cells and FOP knockdown cells were comparable (Figure 2B and 2C). However, the FOP knockdown cells formed significantly longer cilia, compared to the negative control cells (Figure 2B and 2D).

**Figure 2.**
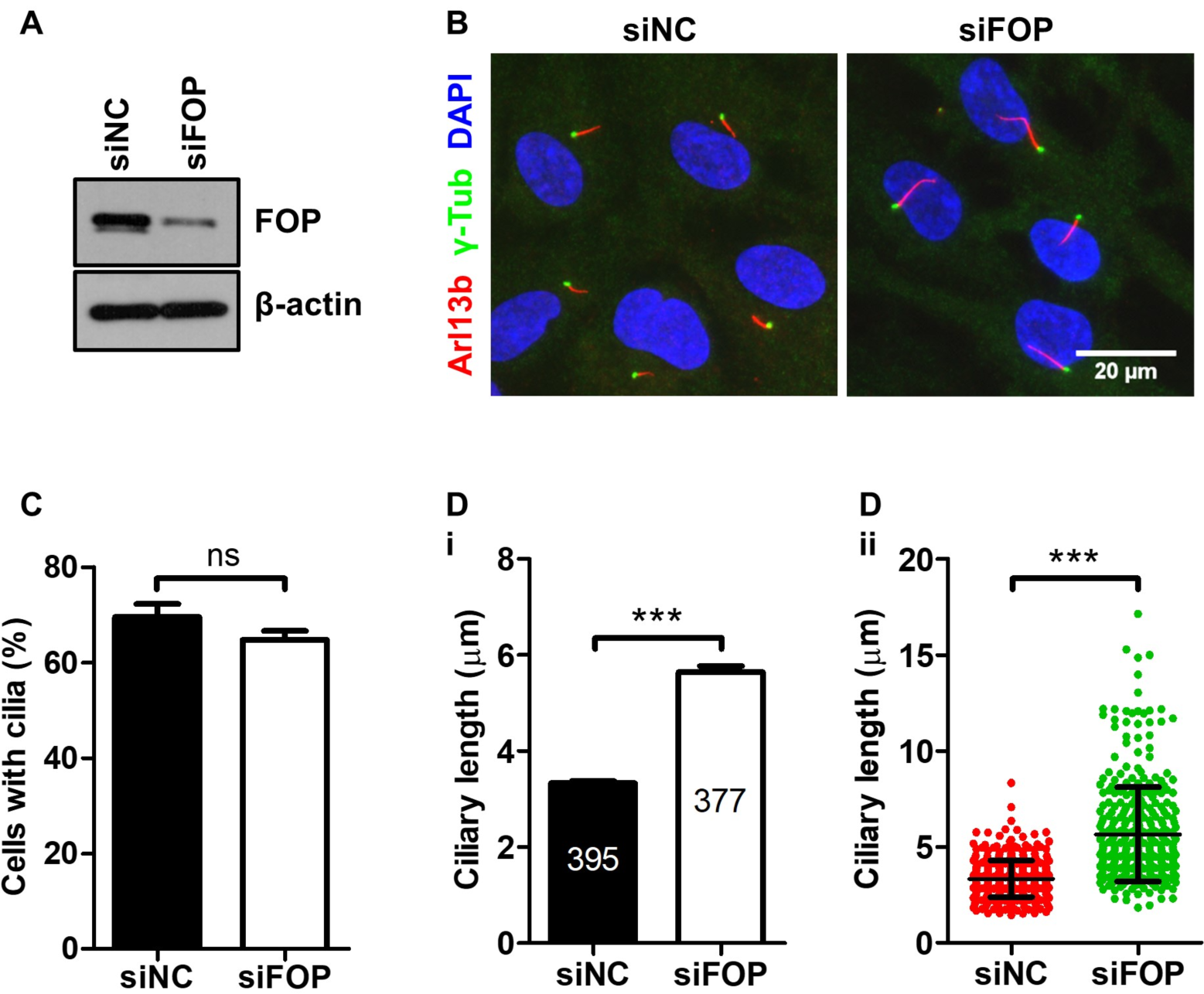
Knockdown of FOP increases ciliary length. (**A**) RPE1 cells were transfected with negative control siRNA (siNC) or FOP siRNA (siFOP). The knockdown efficiency was determined by immunoblotting at 72 hrs post transfection. (**B**) RPE1 cells transfected with negative control siRNA or FOP siRNA were serum-starved and immunostained for Arl13b (red) and *γ*-tubulin (*γ*-Tub; green). The nuclei were stained with DAPI. (**C**) Quantification of the percentage of ciliated cells described in (**B**). At least 200 cells were analyzed for each sample per experiment. Data are presented as mean ± SEM from four independent experiments; ns: not significant; p>0.05 (unpaired, two-tailed, Student’s t-test). (**D, i**) Quantification of the average ciliary length in negative control cells (siNC) and FOP knockdown cells (siFOP). The numbers of cilia measured are indicated in the bars. Data are presented as mean ± SEM; ***p<0.001 (unpaired, two-tailed, Student’s t-test). (**D, ii**) Scatter plot showing individual ciliary length. Data are presented as mean ± SD; ***p<0.001 (unpaired, two-tailed Student’s t-test).

To collaborate these results, we examined the effects of FOP overexpression on primary cilia in a cell line stably expressing FOP-GFP. Overexpression of FOP (Figure 3A) led to a decrease in the percentage of ciliated cells, while the length of the respective cilia also decreased (Figure 3B-D). Together, the data from FOP knockdown and overexpression suggest that FOP suppresses primary cilia formation and elongation.

**Figure 3.**
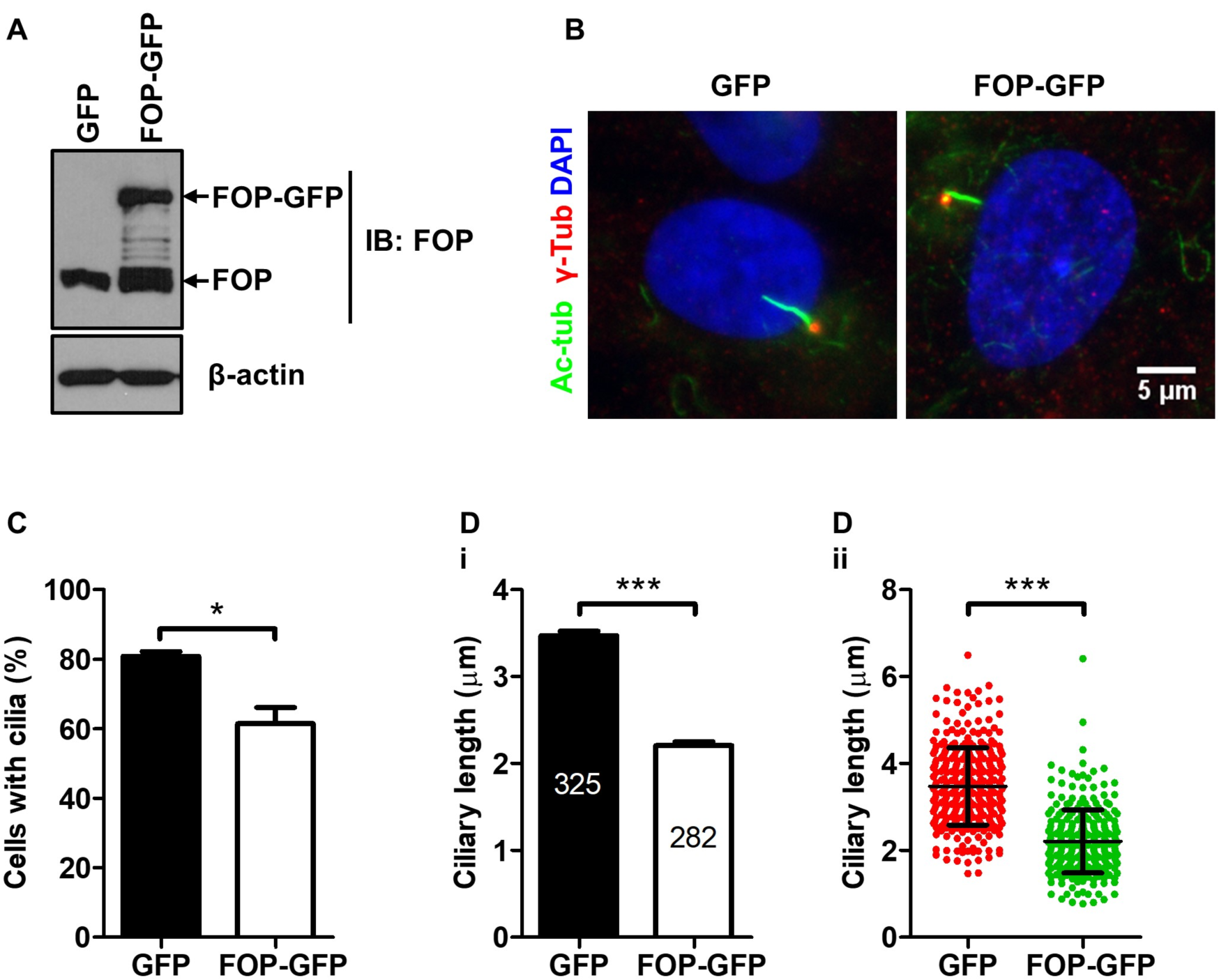
Overexpression of FOP suppresses ciliogenesis. (**A**) Expression of RPE1 cells stably expressing GFP-tagged human FOP. Lysates of cells stably expressing GFP or FOP-GFP were analyzed by immunoblotting with antibodies against FOP. (**B**) Representative immunofluorescence images of the serum-starved vector control cells (GFP) and FOP-overexpressed cells (FOP-GFP) immunostained for acetylated tubulin (Ac-tub; green) and *γ*-tubulin (*γ*-Tub; red). The nuclei were stained with DAPI. (**C**) Quantification of the percentage of ciliated cells. At least 200 cells were analyzed for each sample per experiment. Data are presented as mean ± SEM from three independent experiments; *p<0.05 (unpaired Student’s t-test). (**D, i**) Quantification of the average ciliary length in vector control cells and FOP-overexpressed cells. The numbers of cilia measured are indicated in the bars. Data are presented as mean ± SEM; ***p<0.001 (unpaired, two-tailed, Student’s t-test). (**D, ii**) Scatter plot showing individual ciliary length. Data are presented as mean ± SD; ***p<0.001 (unpaired, two-tailed, Student’s t-test).

As primary cilia are assembled in quiescent cells (when cells exited the cell cycle) (Sánchez and Dynlacht, 2016), and the involvement of FOP in the regulation of cell cycle progression has been previously reported (Acquaviva et al., 2009), we therefore determined if FOP may indirectly inhibit cilia formation and elongation by preventing cell cycle exit. We examined the cell cycle profiles of FOP-depleted and FOP-overexpressed cells after serum starvation, and found that both treatments had little effect on the cell cycle profile (Figure S1A-D). Therefore, the negative effects of FOP on primary cilia did not result from its roles in cell cycle regulation.

As the autophagy-mediated degradation of OFD1 (a homolog of FOP) promotes ciliogenesis (Tang et al., 2013), we determined if knockdown of FOP may promote ciliogenesis by enhancing OFD1 degradation upon serum starvation. OFD1 at the ciliary base was examined by immunostaining. Knockdown of FOP did not significantly affect the degradation of OFD1 after serum starvation (Figure S1E). Therefore, cilia elongation induced by FOP knockdown is not mediated through OFD1 degradation.

### FOP-Promoted Cilia Disassembly Is Dependent on Actin Polymerization

The actin cytoskeleton has been established as a negative regulator of ciliogenesis (Bershteyn et al., 2010; Drummond et al., 2018; Sharma et al., 2011). Disruption of the actin cytoskeleton structure with Cytochalasin D (CytoD), an inhibitor of actin polymerization, induces primary cilia formation and elongation (Sharma et al., 2011). Moreover, depletion of proteins that are required for actin nucleation, branching, polymerization, or stress fiber formation also promotes ciliogenesis (Cao et al., 2012; Drummond et al., 2018). We therefore determined if FOP promotes cilia shortening by strengthening the actin cytoskeleton. We observed that FOP-overexpressed cells (FOP-GFP) displayed increased F-actin staining and stress fibers, accompanying a lower frequency of ciliation and shorter cilia (Figure 4A-C), suggesting that the negative role of FOP in ciliogenesis may be correlated with its positive role in actin cytoskeleton formation.

**Figure 4.**
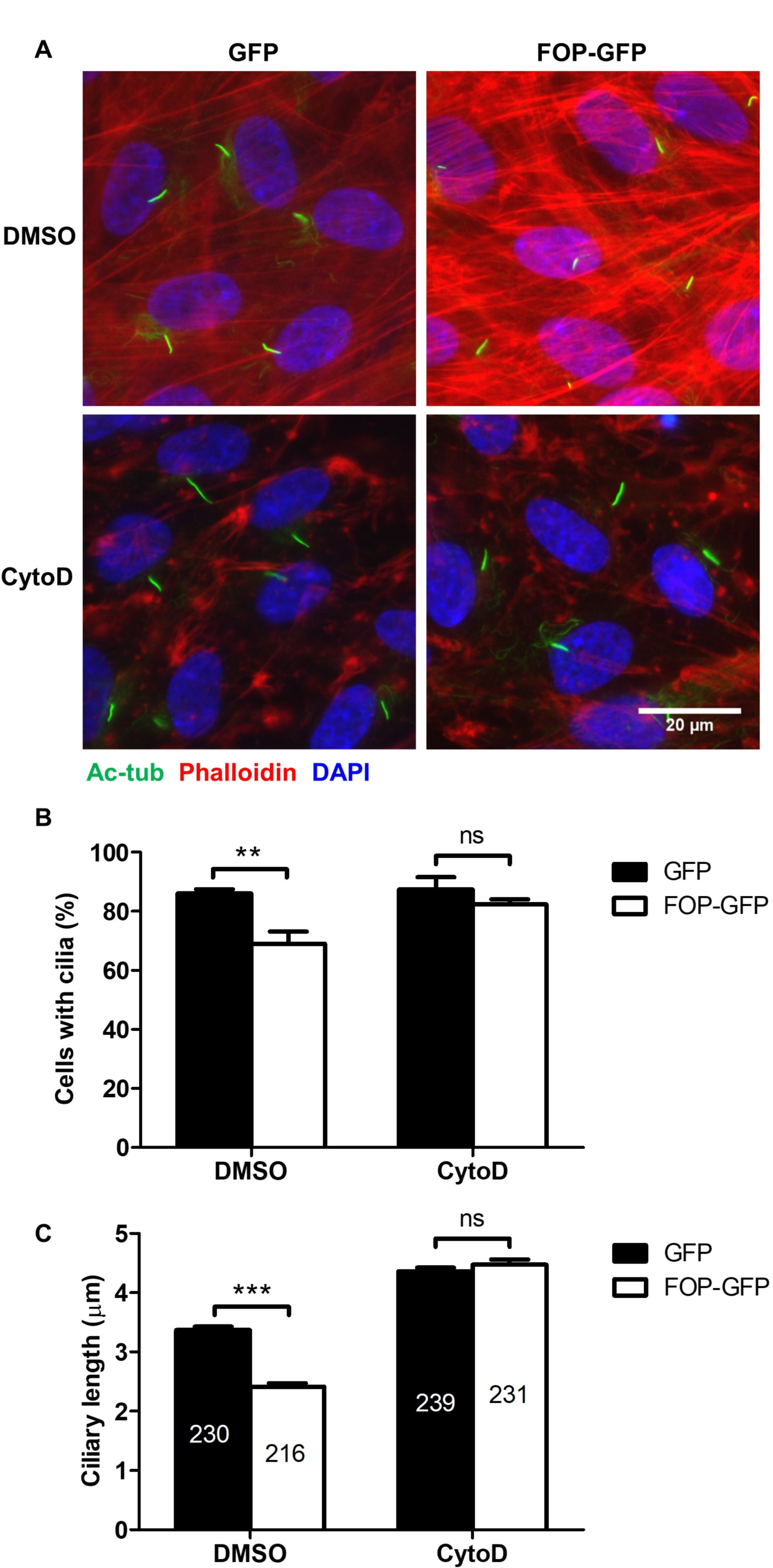
Inhibition of actin polymerization with Cytochalasin D abrogates FOP-promoted cilia shortening. (**A**) RPE1 cells stably expressing GFP or FOP-GFP were serum-starved for 48 hrs and treated with Cytochalasin D (CytoD) for 2 hrs prior to fixation. Fixed cells were immunostained for acetylated *α*-tubulin (Ac-tub; green) and Phalloidin (red). The nuclei were stained with DAPI. (**B**) Quantification of the percentage of ciliated cells described in (**A**), at least 200 cells were analyzed for each sample per experiment. (**C**) Quantification of the average ciliary length. The numbers of cilia measured are indicated in the bars. Data are presented as mean ± SEM from three independent experiments; **p<0.01; ***p<0.001; ns, not significant (two-way ANOVA followed by Bonferroni’s test).

We then asked whether disrupting the actin structure with CytoD could attenuate the negative effects of FOP in ciliogenesis. We found that after CytoD treatment, the FOP-overexpressed cells no longer showed a decrease of ciliation or cilia length compared to the vector control cells, (Figure 4A-C) indicating that the CytoD treatment negated the negative effects of FOP overexpression on ciliogenesis. Therefore, these data suggest that FOP promotes cilia shortening through enhancing actin polymerization.

### FOP Is Required for Timely Cilia Disassembly during Cell Cycle Re-entry

Since FOP negatively regulates cilia length, we next determined whether FOP promotes cilia disassembly when quiescent cells re-enter the cell cycle upon serum re-stimulation. To this end, FOP-silenced RPE1 cells were serum starved to induce cilia formation and then re-stimulated with serum to induce cilia resorption and cell cycle re-entry. Consistent with the data shown in Figure 2, the percentage of ciliated cells in the negative control cells and FOP-silenced cells were similar after serum starvation, while FOP-silenced cells possessed longer cilia (0 hr; Figure 5A-C). In response to serum stimulation, the cilia in the negative control cells were resorbed (Figure 5A-C), as previously reported (Pugacheva et al., 2007). In contrast, this process was suppressed in FOP-silenced cells: at all of the time points after serum addition, FOP-silenced cells had a significantly higher frequency of, and longer cilia than the negative control cells (Figure 5A-4C). It should be noted that, in FOP-silenced cells, the length of primary cilia slightly decreased after 2 hrs of serum re-stimulation, but gradually increased at 6-24 hrs after serum re-stimulation.

**Figure 5.**
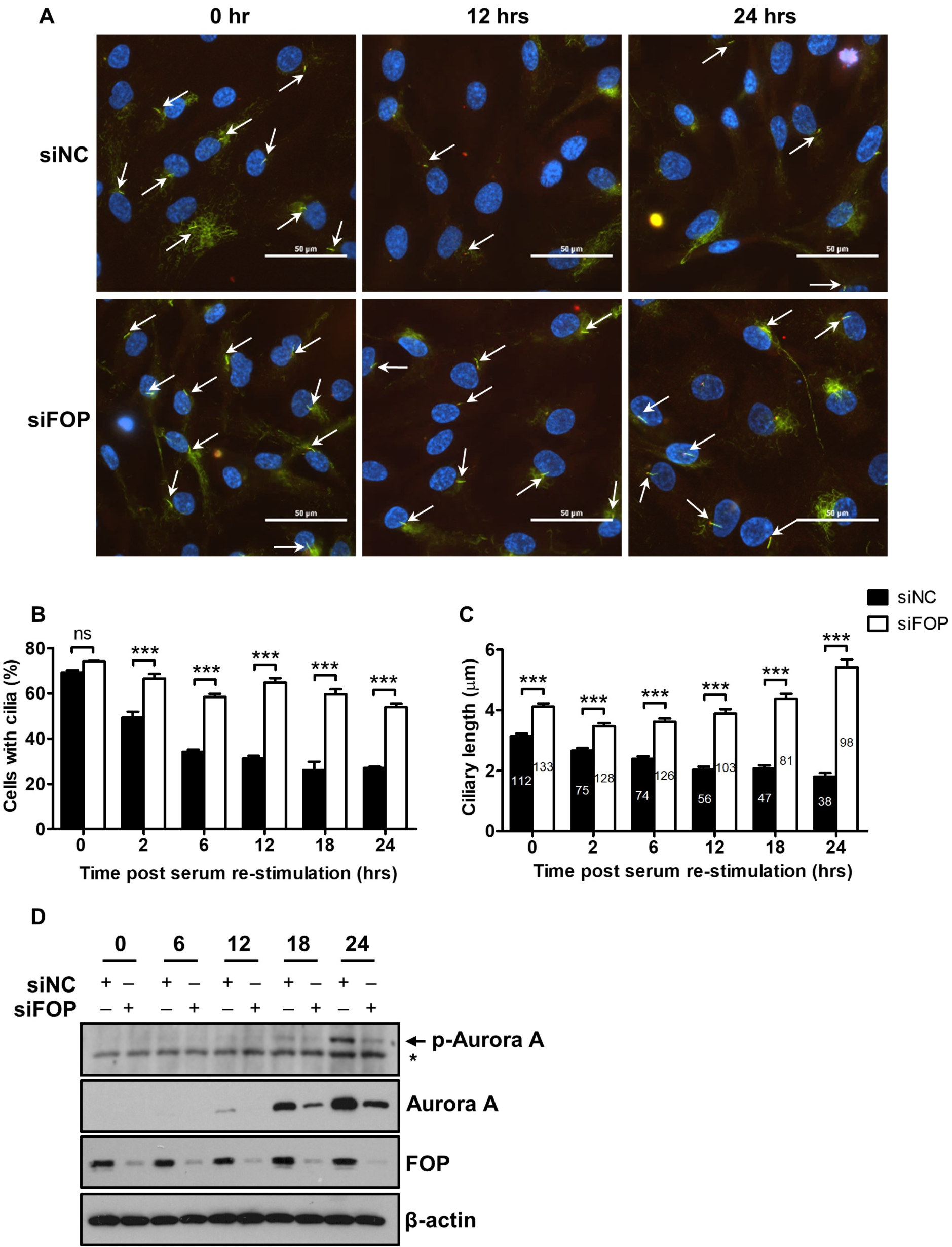
FOP is required for timely cilia disassembly. (**A**) Cells transfected with negative control siRNA or FOP siRNA were serum-starved for 48 hrs, followed by serum re-stimulation and fixation at different time points as indicated in (**B** and **C**). Fixed cells were immunostained for acetylated *α*-tubulin (Ac-tub; green) and *γ*-tubulin (*γ*-Tub; red). Representative photos are shown for the 0 hr, 12 hrs and 24 hrs samples. The nuclei were stained with DAPI. White arrows indicate primary cilia. (**B**) Quantification of the percentage of ciliated cells described in (**A**). At least 200 cells were analyzed for each sample per experiment. (**C**) Average cilia length described in (**A**). The number of cilia measured were indicated in the bars. Data are presented as mean ± SEM from three independent experiments; ***p<0.001; ns, not significant (two-way ANOVA followed by Bonferroni’s test). (**D**) Phosphorylation level of Aurora A at Thr288 (p-Aurora A) and total level of Aurora A in siNC or siFOP treated cells at 0 hr, 6 hrs, 12 hrs, 18 hrs, and 24 hrs post serum re-stimulation.

Next, we asked which signaling pathway is responsible for the role of FOP in timely cilia disassembly. One of well-studied axes involved in cilia disassembly during cell cycle re-entry is Aurora A-HDAC6 (Inoko et al., 2012; Kinzel et al., 2010; Plotnikova et al., 2012; Pugacheva et al., 2007). Aurora A can be activated by HEF1, Pifo, trichoplein and calcium/calmodulin, and activated Aurora A in turn phosphorylates HDAC6 and stimulates its tubulin deacetylation activity, resulting in the destabilization of ciliary axoneme and cilia resorption (Inoko et al., 2012; Kinzel et al., 2010; Plotnikova et al., 2012; Pugacheva et al., 2007). We therefore investigated the relationship between FOP and Aurora A during cilia disassembly and cell cycle re-entry. Interestingly, immunoblotting analysis showed that the expression levels of Aurora A and phosphorylated Aurora A at T288 (p-Aurora A) decreased in FOP-silenced cells, which was most obvious at 18 hrs and 24 hrs post serum re-stimulation (Figure 5D), suggesting that FOP may modulate the expression and activation of Aurora A to promote cilia disassembly.

### Knockdown of FOP Delays Cell Cycle Re-entry

The disassembly of primary cilia is required for cell cycle re-entry (Kim et al., 2011; Li et al., 2011). As knockdown of FOP suppresses cilia disassembly, we therefore asked whether silencing of FOP inhibits cell cycle re-entry. After serum starvation, cell cycle re-entry was induced by re-addition of serum. Cells were labeled with ethynyl-deoxyuridine (EdU) after serum re-stimulation, and the percentage of EdU positive cells was determined. The percentage of EdU positive cells in FOP-silenced cells was significantly lower than that of the negative control cells at 12 hrs, 18 hrs, and 24 hrs after serum re-stimulation (Figure 6A-B). Immunoblotting analysis also showed that FOP-silenced cells had lower levels of the cell proliferation markers, including phosphorylated Rb at Ser807/811, phosphorylated cdc2 at Tyr15 and cyclin A during cell cycle re-entry (Figure 6C). Correspondingly, flow cytometry analysis also revealed a delay in cell cycle progression (Figure 6D). In addition, 24 hrs after serum re-stimulation, ∼12.3% of control cells entered mitosis, as indicated by the nuclear morphology, whereas only ∼3.6% of FOP knockdown cells were mitotic (Figure 6E). Collectively, these data indicate that the knockdown of FOP delayed cell cycle re-entry.

**Figure 6.**
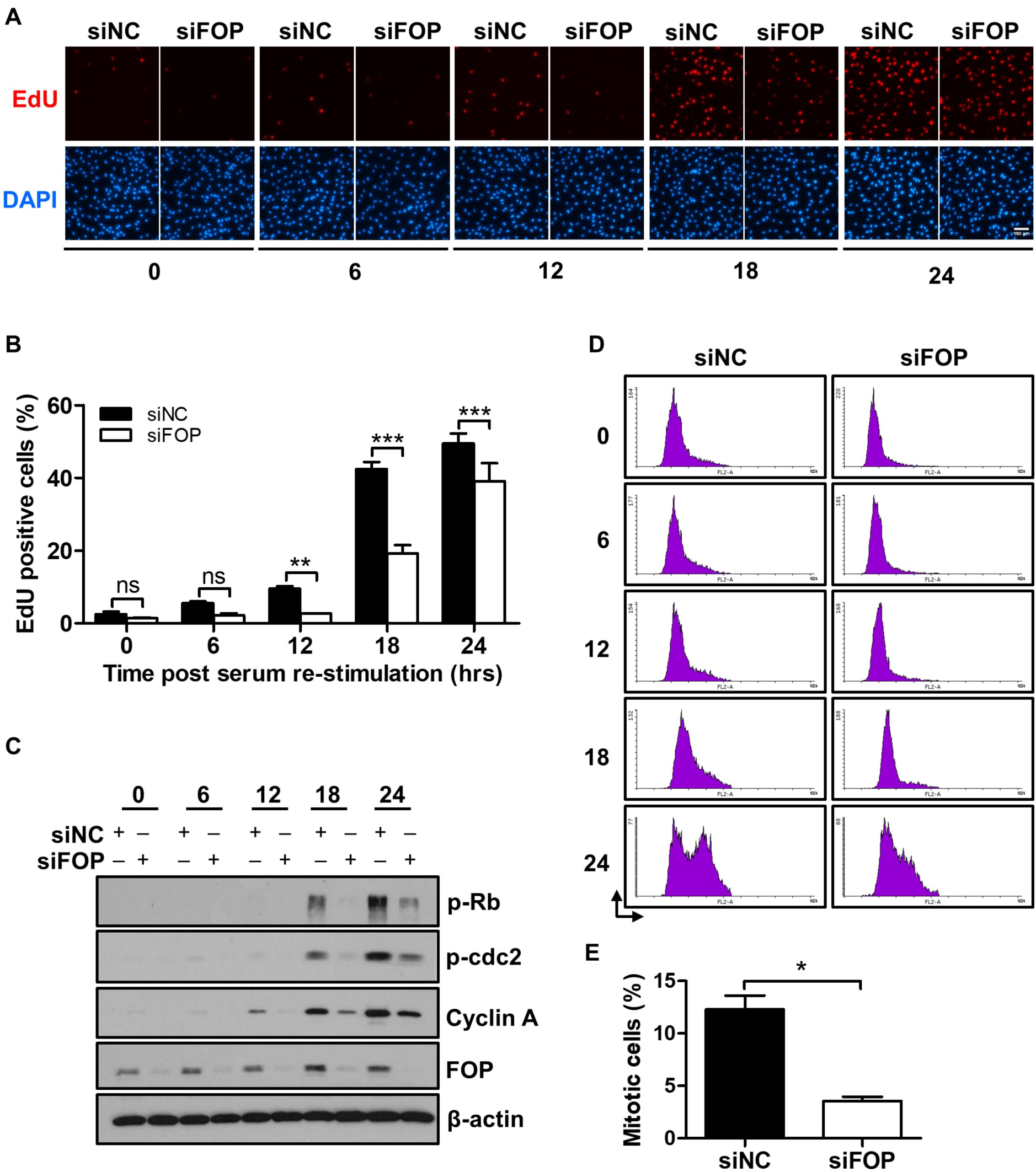
Knockdown of FOP delays cell cycle re-entry. (**A**) RPE1 cells were transfected with control siRNA or FOP siRNA followed by 48 hrs of serum-starvation. Cells were then serum re-stimulated for the indicated durations, being labeled with EdU during the final 2 hrs before cell harvesting. The nuclei were stained with DAPI. (**B**) Quantification of the percentage of EdU positive cells described in (**A**). At least 500 cells were analyzed for each sample per experiment. **p<0.01; ***p<0.001; ns, not significant (two-way ANOVA followed by Bonferroni’s test). (**C**) Phosphorylation levels of Rb at Ser807/811 (p-Rb), cdc2 at Tyr 15 (p-cdc2) and levels of Cyclin A and FOP in siNC and siFOP cells at 0 hr, 6 hrs, 12 hrs, 18 hrs, and 24 hrs post serum re-stimulation. (**D**) Cell cycle profiles of siNC and siFOP treated cells at 0 hr, 6 hrs, 12 hrs, 18 hrs, and 24 hrs after serum re-stimulation were determined by flow cytometry. (**E**) Quantification of the percentage of mitotic cells at 24 hrs after serum re-stimulation. At least 100 cells were analyzed for each sample per experiment. Data are presented as mean ± SEM from three independent experiments; *p<0.05; unpaired, two-tailed (Student’s t-test).

### FOP Regulates Cell Cycle Re-entry through Modulating Primary Cilia

We then asked if the delay in cell cycle re-entry caused by FOP knockdown is mediated by primary cilia. We silenced IFT20, a component of intraflagellar transport, to disrupt primary cilia formation (Follit et al., 2006). Consistent with previous studies, the depletion of IFT20 strongly inhibited cilia assembly (Figure S2A-C). In addition, the primary cilia were also shortened after IFT20 depletion in cells transfected with either negative control siRNA or FOP siRNA (Fig. S2D and E).

We then investigated if the loss of cilia by IFT20 silencing can overcome the delay in cell cycle re-entry induced by FOP depletion. Knockdown of FOP without ITF20 silencing (siNC, which was the control siRNA for the IFT20 siRNA) led to a significant decrease in the percentage of EdU positive cells at 18 hrs after serum re-stimulation, confirming the delay in cell cycle re-entry as described above. In contrast, this delay was rescued by IFT20 depletion (Figure 7A and B). Correspondingly, the levels of phosphorylated Rb (p-Rb), phosphorylated cdc2 (p-cdc2) and Cyclin A in the FOP-silenced cells were also restored by IFT20 depletion (Figure 7C). Flow cytometry analysis data also showed that the inhibition of ciliogenesis by IFT20 depletion rescued the delay of cell cycle progression induced by FOP knockdown (Figure 7D). These results collectively suggest that the delay in cell cycle re-entry induced by FOP knockdown is dependent on primary cilia.

**Figure 7.**
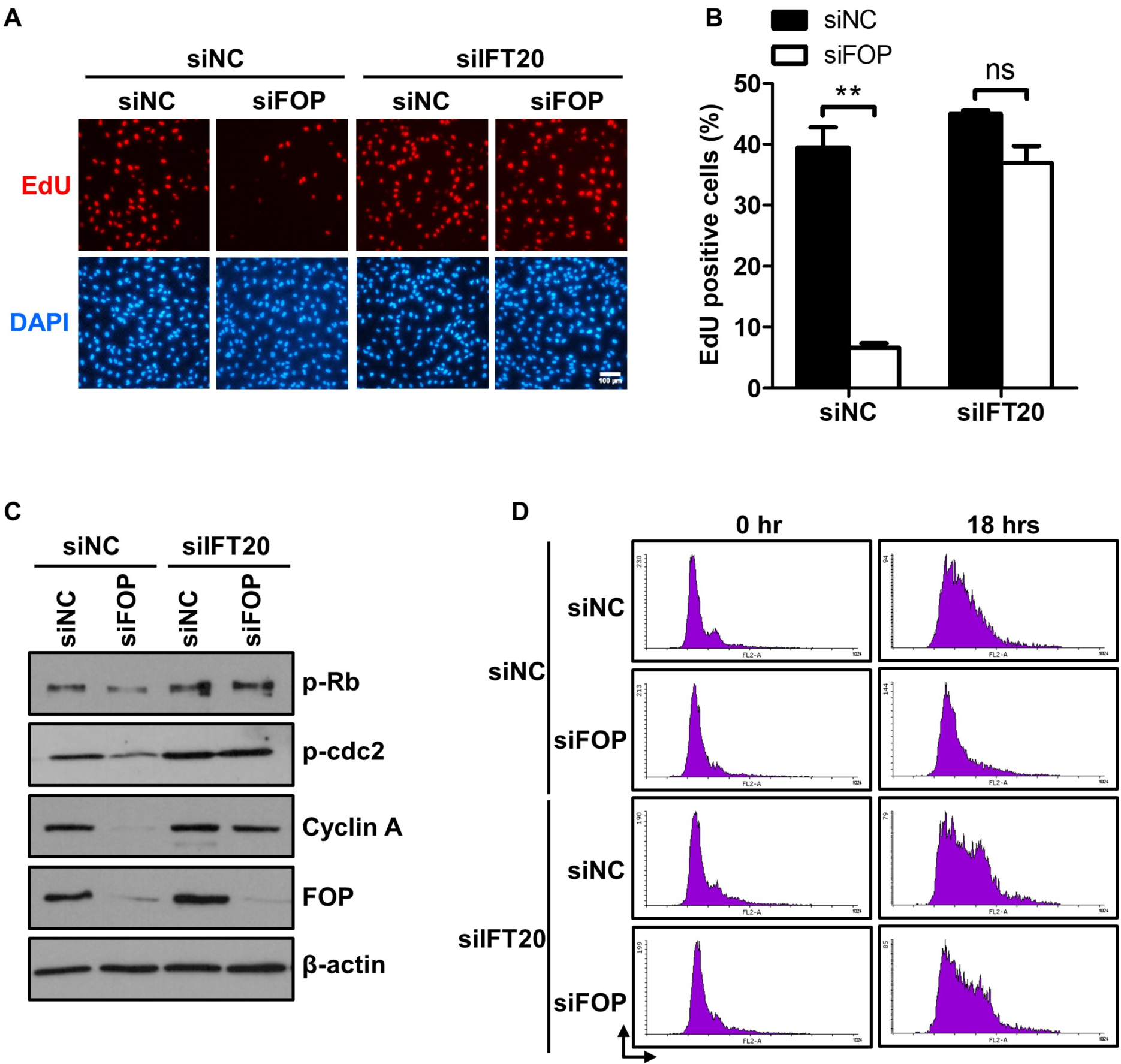
Delay in cell cycle re-entry induced by FOP knockdown is cilia-dependent. (**A**) RPE1 cells were transfected with the indicated siRNAs. Following 48 hrs of serum-starvation, the cells were serum re-stimulated for 18 hrs, being labeled with EdU during the final 2 hrs. The nuclei were stained with DAPI. (**B**) Quantification of the percentage of EdU positive cells described in (**A**). At least 500 cells were analyzed per condition per experiment. Data are presented as mean ± SEM from three independent experiments; **p<0.01; ns, not significant (two-way ANOVA followed by Bonferroni’s test). (**C**) The protein levels of phosphorylated Rb at Ser807/811 (p-Rb), phosphorylated cdc2 at Tyr 15 (p-cdc2), Cyclin A and FOP in cells transfected with indicated siRNA at 18 hrs post serum re-stimulation. (**D**) Cell cycle profiles of cells transfected with the indicated siRNA at 0 hr and 18 hrs post serum re-stimulation were determined by flow cytometry.

## Discussion

Although recent studies reported that near complete silencing of FOP inhibited primary cilia formation (Cabaud et al., 2018; Kanie et al., 2017; Mojarad et al., 2017), others found that moderate knockdown of FOP had no effect on ciliogenesis (Graser et al., 2007; Lee and Stearns, 2013). Here we have demonstrated that FOP plays a negative role in ciliogenesis. First, the expression of FOP decreases as primary cilia are formed upon serum starvation, and gradually increases as primary cilia are disassembled after serum re-stimulation, suggesting that a negative correlation of FOP with ciliogenesis. Secondly, knockdown of FOP increases the length of primary cilia. Thirdly, ectopic expression of FOP suppresses primary cilia formation and elongation. The results from both the knockdown and overexpression experiments minimized the possibility of off-target effects. Lastly, we show that FOP promotes cilia disassembly by enhancing actin cytoskeleton formation.

Expression of the proteins involved in ciliogenesis must be precisely regulated to exert different roles during cilia assembly and disassembly. For example, CP110 displays complex roles in ciliogenesis through the ubiquitin-proteasome system as well as transcriptional programs. High levels of CP110 suppress primary cilia formation, while optimal levels of CP110 promote ciliogenesis (Cao et al., 2012; Kobayashi et al., 2011; Song et al., 2014; Spektor et al., 2007; Walentek et al., 2016; Yadav et al., 2016). Notice that FOP does not completely degrade during ciliogenesis. Therefore, it is possible that a small fraction of FOP is necessary and sufficient for the recruitment of the CEP19-RABL2 complex to the ciliary base, allowing IFT entry and initiation of ciliogenesis. As such, complete absence of FOP would compromise the essential role of FOP in the early steps of primary cilia assembly. In the present study, we identified a novel function of FOP in ciliogenesis. We found that FOP suppresses ciliary axoneme elongation and promotes timely cilia disassembly during cell cycle re-entry, while knockdown of FOP by 80% does not impair primary cilia assembly. Together with the data from others’ previous studies, our results suggest that FOP may play multiple and dose-dependent roles in ciliogenesis, being required for primary cilia formation and also playing a negative role in the regulation of cilia length. It will be interesting to elucidate how FOP’s levels are precisely controlled to produce optimal levels for ciliogenesis and disassembly.

Recently, many studies have suggested that actin dynamics regulate cilia assembly and disassembly (Bershteyn et al., 2010; Cao et al., 2012; Drummond et al., 2018). Treatment with the actin polymerizing inhibitor, Cytochalasin D (CytoD), leads to a higher percentage of ciliated cells and longer primary cilia, while serum-induced cilia resorption is also blocked (Drummond et al., 2018; Kim et al., 2011; Li et al., 2011; Sharma et al., 2011). Many actin regulatory proteins, such as MIM, cortactin and ARP2/3, have been reported to be involved in ciliogenesis (Bershteyn et al., 2010; Cao et al., 2012; Drummond et al., 2018; Saito et al., 2017). Our preliminary proteomic analysis by mass spectrometry (unpublished data, H. Jiang and C. Liang) as well as a previous study (Kazazian et al., 2017) found that several of the FOP-interacting proteins are also actin-associated, two of which being ARP2 and ARP3. The ARP2/3 complex is crucial for branched F-actin formation (Pizarro-Cerdá et al., 2017). It has been proposed that branched F-actin prevents the formation of pericentrosomal recycling endosomes that transport membranes for ciliogenesis. Depletion of either ARP2 or ARP3 promotes cilia formation and cilia elongation (Cao et al., 2012; Drummond et al., 2018). In this study, we show that overexpression of FOP promotes F-actin stress fiber formation, and that pharmacological inhibition of actin polymerization with Cytochalasin D abrogates FOP-induced cilia disassembly. Together, these data suggest that FOP promotes cilia disassembly by enhancing actin cytoskeleton formation.

The Aurora A-HDAC6 signaling pathway is essential for cilia disassembly when cells re-enter the cell cycle (Pugacheva et al., 2007). Aurora A has several activators including HEF1, Pifo and trichoplein (Inoko et al., 2012; Kinzel et al., 2010; Pugacheva et al., 2007). Our data suggest that FOP may serve as an upstream regulator of Aurora A and modulate the expression and activation of Aurora A, thereby promoting cilia disassembly. Although It has previously been suggested that Aurora A-HDAC6 and actin dynamics are two independent pathways that regulate cilia disassembly (Bershteyn et al., 2010), a recent study demonstrated that HDAC6 also deacetylates another substrate, cortactin, which further enhances actin polymerization, thus inducing cilia disassembly (Ran et al., 2015; Zhang et al., 2007). Together with these previous findings, our data on the regulatory role of FOP on Aurora A in the current study further strengthen the connection between the Aurora A-HDAC6 pathway and actin dynamics in the regulation of cilia disassembly.

The link between ciliogenesis and the cell cycle has been well established (Kim et al., 2011; Li et al., 2011; Pugacheva et al., 2007). Primary cilia have to be completely disassembled prior to mitosis, releasing the centrioles to form the mitotic spindle poles. Therefore, the length of primary cilia regulates the duration of the cell cycle. For example, depletion of Nde1 or Tctex-1 accelerates cell cycle progression by triggering timely cilia disassembly (Kim et al., 2011; Li et al., 2011). The role of FOP in cell cycle progression has previously been implicated (Acquaviva et al., 2009). The data we present here suggest that FOP, like Nde1 and Tctex-1, also facilitates cell cycle re-entry by promoting cilia disassembly. Given that FOP is a centrosomal protein, and loss of centrosome integrity causes G_1_-S arrest (Mikule et al., 2007; Yan et al., 2006), it could be argued that the cell cycle delay in FOP knockdown cells originated from a loss of centrosome integrity. However, as the cell cycle re-entry delay induced by FOP knockdown can be rescued by the depletion of IFT20 (which is essential for cilia assembly but not centrosome integrity; Follit et al., 2006), a possible loss of centrosome integrity cannot be the reason for cell cycle re-entry delay induced by FOP knockdown. Furthermore, in chicken DT40 lymphocytes, FOP knockout did not impair centrosome integrity (Acquaviva et al., 2009). By immunostaining *γ*-tubulin, we also did not detect any apparent centrosome defects. Therefore, the delay in cell cycle re-entry induced by FOP knockdown is mediated by primary cilia.

Recent studies have suggested that primary cilia are involved in tumorigenesis and tumor progression, as the loss of cilia is frequently observed in various types of cancer such as breast cancer, prostate cancer and pancreatic ductal adenocarcinoma (Hassounah et al., 2013; Menzl et al., 2014; Seeley et al., 2009; Yuan et al., 2010). Although the mechanism is still unclear, the loss of cilia probably provides a growth advantage and promotes malignant transformation and metastasis during the early stages of tumor development (Basten and Giles, 2013; Deng et al., 2018; Zingg et al., 2018). Upregulation of FOP has been observed in lung cancer tissues and cell lines (Mano et al., 2007). Here, we demonstrate that FOP promotes cilia disassembly and accelerates cell cycle progression. It is likely that the upregulation of FOP induces cilia loss and provides growth advantages for cancer cells.

## Acknowledgments

We thank Prof. Robert Zong Qi (HKUST) for providing hTERT-RPE1 cell line, Prof. Karl Herrup (HKUST) for providing the p-Rb antibody, Prof. Randy Y.C. Poon (HKUST) for providing the p-cdc2 antibody, Prof. Yu Li (Harbin Institute of Technology) for providing pEGFP-N1 vector, Dr. Shanshan Liu (Shenzhen University) for providing Alexa Fluor 594 phalloidin and Kelvin K.L. Sou for technical assistance. This work was supported by grants from Foshan Science and Technology Bureau, Guangdong, China (2015IT100132) and Guangzhou Committee of Science and Information Innovation, China (201604020038).

